# Pharmacokinetic properties of Ogerin and its regulation of post-ischemia signaling in mice

**DOI:** 10.64898/2026.10.02.754522

**Authors:** Youfang Zhou, Shambhu Joshi, Xiang-ming Zha

## Abstract

Ogerin, a positive regulator of GPR68, reduces infarct volume and attenuates blood-brain barrier (BBB) leakage post transient middle cerebral artery occlusion (tMCAO). These data support the potential use of Ogerin for ischemic protection. For therapeutic interventions, tissue distribution, kinetics, and brain penetration are important characteristics for a compound. For these reasons, we performed liquid chromatography-mass spectrometry (LC-MS) analysis of Ogerin in mice. At 3 hours following intraperitoneal (IP) injection, liver exhibited highest level of Ogerin among all major organs assessed. We further analyzed time-dependent changes in Ogerin concentration in serum following intravenous delivery. Brain Ogerin level was ∼140 ng/g at 2 hour and ∼0.06ng/g at 24 hours. The brain-to-serum ratio of Ogerin was ∼0.6 at 5 hr. To determine the effect of Ogerin on signaling, we collected samples 24 hours after tMCAO with or without Ogerin delivered and performed Western blotting. Consistent with the diminished level of Ogerin in brain at 24 hour, tMCAO increased similar changes in phosphorylation of PKA substrates and PKC substrates in vehicle- and Ogerin-treated groups. The pharmacokinetic data suggests that Ogerin is effective mainly during acute hours after injection. This finding generates important insights on the design of effective protocol for Ogerin delivery to potentiate GPR68 function in vivo.

## Introduction

Stroke is one of the leading causes of death and long-term disability worldwide [1–3]. Thus, better therapy and new alternative medicines are urgently needed. The most common type of stroke is ischemic stroke [3–5]. Following ischemia, acidosis occurs due to accumulation of CO2 and metabolic disruption [6–8]. During ischemic stroke, pH declines most profoundly (approximately 6.0) in the core area while in the penumbra surrounding the core is around 6.70. Following reperfusion, a second phase of acidosis occurs and can last for at least hours after reperfusion [9–11]. In ischemia, the degree of acidosis positively correlates with the severity of injury [8, 12]. Since acidosis parallels with injury, targeting acid-sensitive pathways would be a good approach to treat strokes.

There are multiple acid sensitive receptors in the mammalian brain, including acid-sensing ion channels (ASICs), proton-activated chloride channel, G-protein coupled receptor 4 (GPR4), G-protein coupled receptor 65 (GPR65), and G protein-coupled receptor 68 (GPR68) [8, 13]. Among these, GPR68 exhibits wide expression in brain neurons [14] and its activation leads to neuroprotection in acidotic and ischemic conditions [8, 14]. GPR68 deletion worsens acidotic and ischemic neuronal injury in vitro and in vivo [15]. Conversely, overexpression of GPR68 suppressed inflammation and reduced ischemic brain injury [14]. In recent studies, we and others have assessed the effect of Ogerin, a positive allosteric modulator of GPR68 [16, 17], in brain ischemia. Ogerin treatment reduces brain injury at 24 hours post stroke. Meanwhile, Ogerin acted protection effects at delayed time point after stroke (3 days post-stroke), it significantly attenuated blood-brain barrier (BBB) leakage, brain infarct, and brain edema [14, 15, 18, 19]. These results indicated that Ogerin is a promising drug for improving ischemia outcome.

For better drug targeting in vivo, a better understanding of PKPD of a compound is critical. However, this information is lacking in the literature for Ogerin. In addition, our previous study showed that part of Ogerin-induced protection is through the activation of PERK branch of unfolded proteins responses (UPR) during 3-6 hours of ischemia-reperfusion [19]. How Ogerin alters signaling at later time points of reperfusion remains unclear. Here we performed LC-MS analysis of Ogerin in brain, serum, and major organs at different time points after delivery. We performed time-dependent characterization of Ogerin level in serum to determine its kinetics. Lastly, to better understand the impact of signaling by Ogerin, we performed Western blotting analysis.

## Material and methods

### Mice

Breeding colonies of GPR68+/+ (WT) and GPR68-/-(KO) C57BL/6 mice were maintained as breeding colonies at Tulane University. Both male and female mice were used in this study. All animals were housed under standard laboratory conditions and followed ARRIVE and STAIR guidelines for *in vivo* investigations [20, 21]. All procedures were conducted in accordance with the National Institutes of Health Guide for the Care and Use of Laboratory Animals and were approved by the Tulane University Institutional Animal Care and Use Committee.

### Transient middle cerebral artery occlusion (tMCAO)

tMCAO surgery was performed similarly as described in our earlier studies [18, 19]. Briefly, we used the Longa Method to induce occlusion [22]. Male and female mice were randomly assigned to experimental groups. Anesthesia was induced using isoflurane (1.5%) inhalation in a mixture of oxygen and nitrous oxide (30%/70%). Body temperature was maintained at 37 ± 0.5 °C using a feedback-controlled heating pad. Buprenex (0.05-0.2 mg/kg) was given per-surgery to provide analgesia and alleviate any pain and discomfort. The cerebral blood flow (CBF) was monitored using laser Doppler flowmetry (MoorVMS-LDF2) continuously. A silicone-coated monofilament was inserted via the external carotid artery and carefully advanced into the internal carotid artery to occlude the origin of the middle cerebral artery, and the CBF dropped by at least 80% from the peak value prior to occlusion. After 60 minutes of occlusion, the filament was gently removed to restore blood flow and confirmed by reperfusion. Animals were placed in a clean warmed cage to recover after surgery.

### Ogerin delivery and tissue collection

Ogerin (sigma, SML1482) was dissolved in DMSO (USP grade, UFC Biotechnology #D67685-995) to make 20 mM stock solution. 0.9% NaCL (USP grade, Lab Alley) was used to dilute the stock to working solution (280 μl 0.9% NaCL + 20 μl 20 mM Ogerin). 3 mg/kg of Ogerin was delivered by retro-orbital intravenous (IV), and 10 mg/kg via intraperitoneal (IP). Ogerin was delivered at reperfusion (Figure 3, 5) or 2 hours after reperfusion (Figure 1, 2, 4).

**Figure 1.**
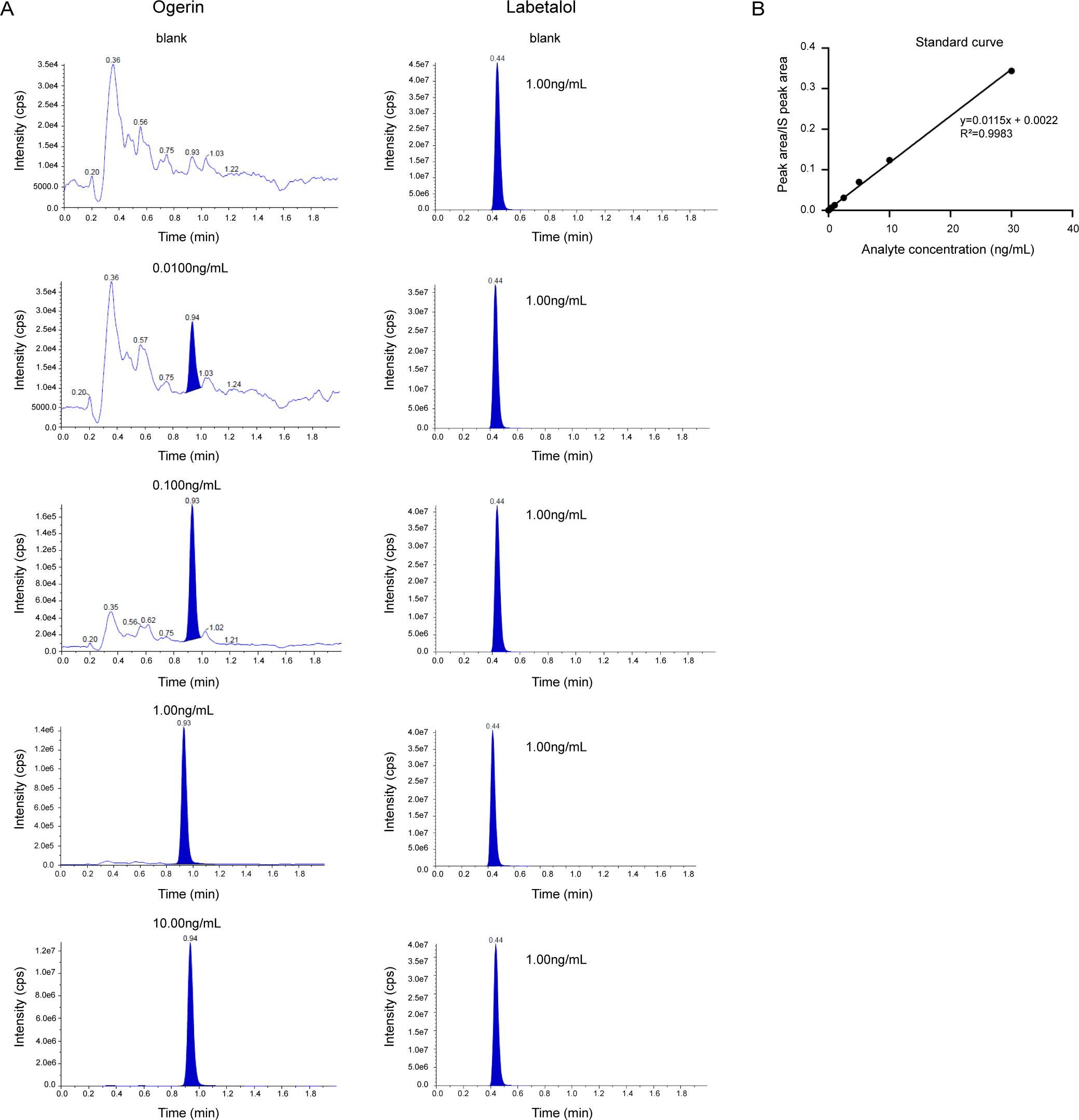
Chromatograms and standard curve. (A) Representative chromatograms of Ogerin (left column) and labetalol (right column). (B) Standard curve of Ogerin.

### Blood and tissue collection

After 3 mg/kg of Ogerin was delivered IV, we collected submandibular blood at 5, 15, 30, 60 min (each time ∼50μl). At 120 min designed endpoint, we collected blood from heart. The collected blood was sitting at room temperature around 30 minutes, then centrifuge at 5000 rpm for 10 minutes and collect the supernatant serum and store at -80°C. For some animals, we collected submandibular blood before Ogerin delivery and centrifuged for serum as pre-serum. Following the terminal blood collection, we quickly dissected liver, kidney, lung, spleen, heart and brain. For brain, we removed cerebellum and brain stem, and separated brain from middle for contralateral and ipsilateral parts. All tissues were collected as fast as possible and stored at -80°C until analysis.

### LC-MS analysis

LC-MS analysis was performed by WuXi AppTec through a service contract. For Serum samples, 40 μl serum was protein-precipitated with 280 µL IS solution (100 ng/mL Labetalol & 100 ng/mL Tolbutamide in 80% MeOH), the mixture was vortex-mixed well and centrifuged at 12,000 rpm for 15 min under 4°C. 200µL supernatant was transferred to a 96-well plate and blow-dried using N2 gas at 35°C. Then the plate was reconsitituted using 125 μL of 10% Acetonitrile in water with 0.1% Formic Acid, then shaken at 800 rpm for 10 min.

For solid tissue samples, tissue samples were homogenized in a ratio of 1:5 (wt/vol) using homogenate buffer (1x PBS: MeOH, v/v = 2:1). 100µL homogenate sample was protein-precipitated with 700 µL IS solution (100 ng/mL Labetalol & 100 ng/mL Tolbutamide in 80% MeOH), the mixture was vortex-mixed well and centrifuged at 12,000 rpm for 15 min, 4°C. 600µL supernatant was transferred to a 96-well plate and blow-dried using N2 gas at 35°C. Then the plate was reconsitituted using 125 µL of 10% Acetonitrile in water with 0.1% Formic Acid, then shaken at 800 rpm for 10 min. Then 3 µL supernatant was injected and on to LC column and analyzed by a Triple Quad 6500+.

### Western blot

Brain samples were lysed in SDS lysis buffer (1%Triton+ 0.4%SDS+ proteinase inhibitor+ phosphatase inhibitor + 1xPBS), homogenized and sonicated, followed by centrifuging at 16000g for 10min at 4°C and collected supernatant. Protein concentration was analyzed using Bio-Rad protein assay. Protein samples were separated with 10% gels and transferred to membranes, with 1 hour blocking (0.1 % casein, 0.2 x PBS, pH 7.2-7.6) at room temperature. Then incubated in primary antibodies at 4°C, overnight. On the next day, the membranes were washed with PBST for 3 times (each 10 minutes), followed by 1 hour incubation with secondary antibodies. After washing 3 times with PBST, membranes were imaged using a Bio-Rad ChemiDoc MP Imaging System. Primary antibodies were used: pPKCSS (CST#2261S, 1:1000), pPKASS (CST#9624S, 1:1,500), pCREB (CST#9198L, 1:2000), CREB (PTG lab# 67927-1-Ig, 1:1,000), ATF6 (Proteintech#24169-1-AP, 1:1,000), IRE1 (CST#3294S, 1:1,000), GRP78/BIP (Proteintech#11587-1-AP, 1:1,000), mTOR (Proteintech#28273-I-AP, 1:2,000), pmTOR (Proteintech#67778-1-Ig, 1:2,000), pJNK (Sigma#07-175, 1:1,000), JNK1 (CST#3708S, 1:2,000), tubulin (DSHB#E7-C, 1:10,000). Secondary antibodies were used: Donkey anti-rabbit IRDye 800CW (Li-Cor # 926-32213, 1:10,000), Donkey anti-mouse IRDye 680LT (Li-Cor#926-68022, 1:10,000).

### Statistical analysis

We performed one-way ANOVA analysis with Tukey for multiple groups comparison, and T-test or Mann-Whitney U test for two groups comparison. All data are presented as mean ±SEM (standard error of the mean) and analyses were performed using GraphPad Prism.

## Results

### Tissue distribution of Ogerin after intraperitoneal injection

To determine the distribution of Ogerin in vivo, we performed Liquid chromatography– mass spectrometry (LC-MS) analysis. For calibration, a standard calibration curve was generated first. Ogerin was added into lysate or serum at was 0.00 (blank), 0.01, 0.02, 0.05 ng/mL, 0.1, 0.2, 1, 5, 10 ng/mL. 1 ng/mL labetalol (IS) was added as internal calibration control. Figure 1A shows a set of representative chromatograms. From the chromatogram readings, we generated the standard curve for calibrating the values from in vivo samples (Figure 1B).

Previous studies have assessed intraperitoneal (IP) delivery of 10 mg/kg Ogerin [23]. To determine tissue distribution of Ogerin, we performed IP of Ogerin in sham-operated C57BL6 mice and collected major organs at 3 hours after delivery, and analyzed Ogerin level by LC-MS. Ogerin exhibited highest level in liver with an average concentration of 208.33 ± 64.09 ng/g (Mean±SEM). Ogerin level in other organs were lower: 22.41ng/g (mean) in heart, 24.81 ng/g in spleen, 38.50 ng/g in lung, and 46.43 ng/g in kidney (Figure 2A). Next, we analyzed Ogerin concentration in serum and brain samples following 60 minutes tMCAO and IP delivery of 10 mg/kg Ogerin. The results showed that the concentration of Ogerin in serum in tMCAO group is ∼38.10 ng/mL in average and ∼16.77 ng/mL in average in sham group (Figure 2B). Ogerin levels in the brain were lower, 11.71 ng/g in sham. Following tMCAO, Ogerin in the brain was 4.28 & 3.43 ng/g in contralateral and ipsilateral side of the post-MCAO brain, respectively (Figure 2C). We analyzed the average brain to serum ratio, for samples which have both measurements available. The average ratio were 0.65 in sham and 0.62 in contralateral and 0.50 in ipsilateral in tMCAO samples.

**Figure 2.**
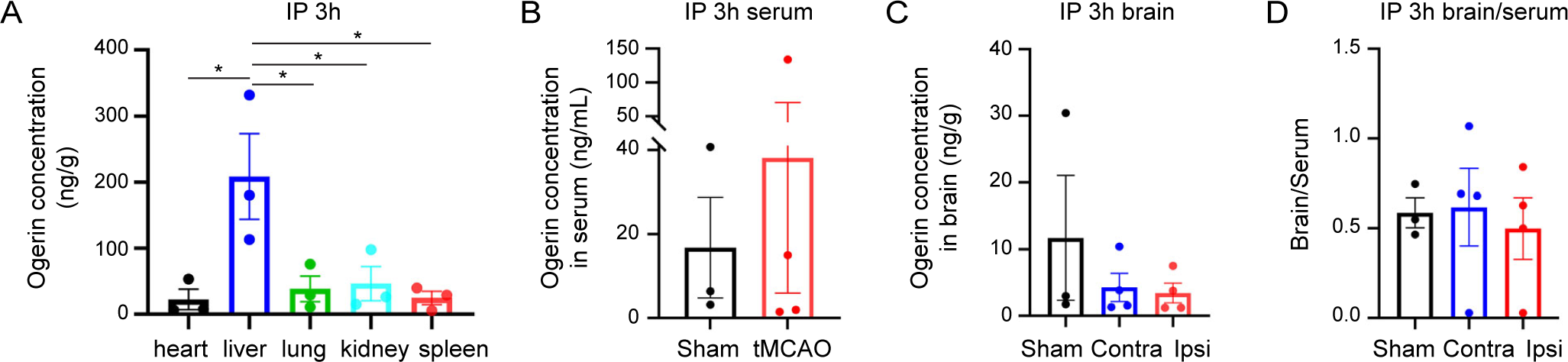
Ogerin distribution in tissues and serum. (A) Ogerin concentration (ng/g) in heart, liver, lung, kidney and spleen in sham mice. (B) Ogerin concentration (ng/mL) in serum in sham or tCMAO mice. (C) Ogerin concentration (ng/g) in brain in sham or tMCAO (contralateral and ipsilateral) mice. (D) brain-serum ratio of sham and tMCAO mice. Samples were collected after 3 hours of IP delivery of Ogerin (10mg/kg) in sham (n=3, female) or tMCAO (n=4, female) mice. Ogerin was delivered 2 hours after reperfusion in tMCAO mice.

### Pharmacokinetics of Ogerin in serum following intravenous delivery

In our previous studies, we performed intravenous (IV) delivery of Ogerin which reduced brain infarct after tMCAO [18, 19]. To better understand the pharmacokinetics of Ogerin in vivo, we delivered Ogerin at 3 mg/kg via IV and collected serum at 5, 15, 30, 60 and 120 minutes from sham and 60-minute tMCAO mice. At 5 minutes after IV injection, the Ogerin concentration in serum was around 1.97 μg/mL, followed by exponential decay from 15 to 120 minutes (Figure 3A). At 120 minutes, the average concentration of Ogerin was around 288.5 ng/mL (Figure 3A). To better calculate the elimination constant, we normalized the Ogerin level in each animal to its 5-min level, and plotted the percentage of Ogerin in serum at different time points. The curve fit a two-component decay with an alpha decay (α) of around 11 minutes and beta (elimination half-life, β) of around 66 minutes (Figure 3B). We also analyzed the Ogerin concentration in the brain at 2-hour end point, and found that at this time, Ogerin in brain was 157.35 and 122.68 ng/g in right and left sides, respectively (Figure 3C). We calculated the average brain-to-serum ratio, for samples that had both measurements available. The average ratio were 0.79 in right side and 0.54 in left side.

**Figure 3.**
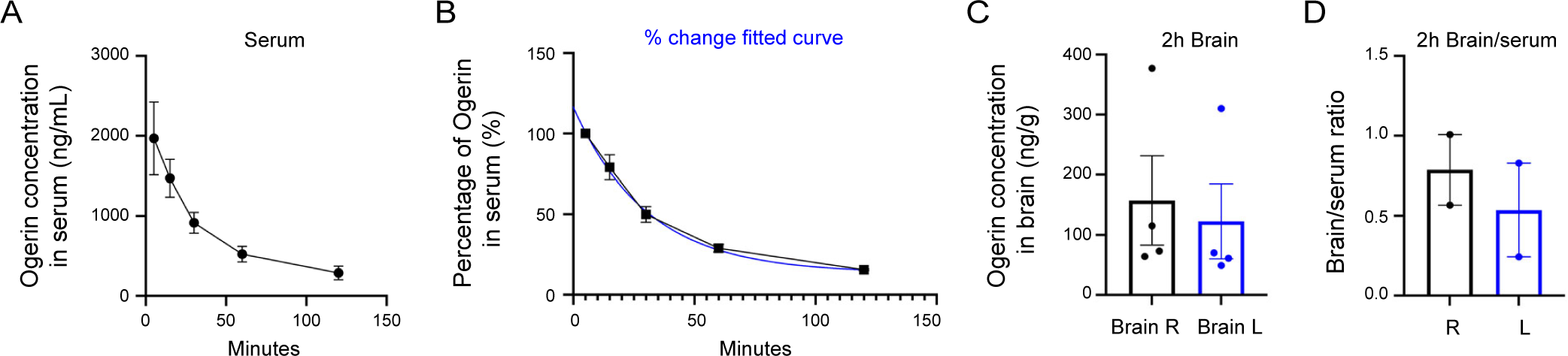
Ogerin dynamics in serum. (A) Ogerin concentration (ng/mL) in serum at 5, 15, 30, 60,120 minutes. (B) fitted curve (blue) based on percentage of Ogerin (%) in serum (black, normalized to Ogerin concentration at 5 minutes). (C) Ogerin concentration (ng/g) in brain at 2 hours. (D) Brain-serum ration at 2 hours end point. 3mg/kg of Ogerin was delivered via IV in sham or tMCAO (at reperfusion) mice. Animals used here are female mice, n=3 sham+1 tMCAO, except at 60 and 120 minutes (female, n=3 sham+1 tMCAO)

Next, we analyzed the Ogerin level at 24 hours after stroke. We performed 60 minutes of tMCAO and delivered 3 mg/kg Ogerin via IV 2 hours after reperfusion, and collected brain and serum samples 24 hours after reperfusion. The results showed that ∼24 hours after IV delivery, the Ogerin level was low in serum (∼0.02 ng/mL) and below detection limit in brain in the sham group (Figure 4A, B). In the tMCAO group, 24h serum showed ∼0.23 ng/mL Ogerin, and the brain showed ∼0.11 ng/g in contralateral and 0.06ng/g in ipsilateral side (Figure 4A, B). As expected, with a second dose of Ogerin at ∼20 hours after stroke, the concentration of Ogerin in serum was ∼4.73 ng/mL, and ∼0.7 ng/g in the brain (0.72 and 0.66 ng/g in contralateral and ipsilateral hemispheres, respectively) (Figure 4A, B).

**Figure 4.**
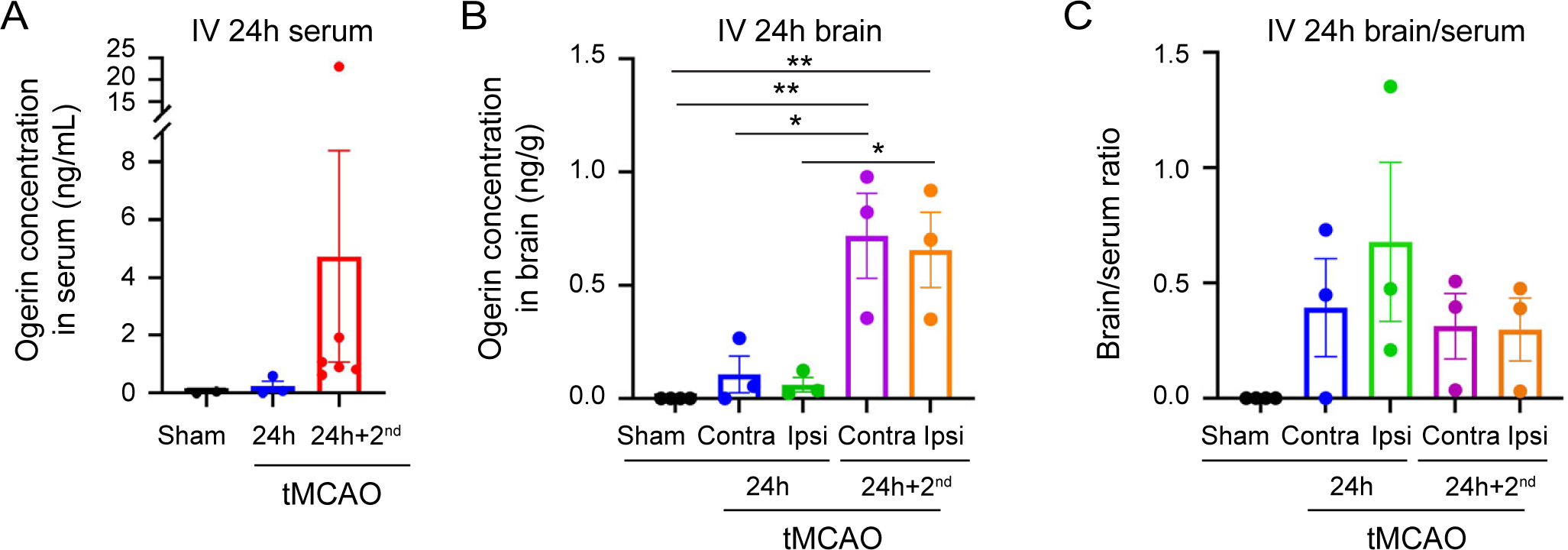
Ogerin decreased rapidly in 24 hours and a second dose helps maintain a high concentration. (A) Ogerin concentration (ng/mL) in serum 24 hours after delivery or a second dose in sham (n=2) and tMCAO (n=3 at 24 hours, n=6 with a second does) mice. (B) Ogerin concentration (ng/g) in brain samples 24 hours after delivery or with a second dose of Ogerin in sham (n=4) and tMCAO (n=3 at 24 hours, n=3 with a second does) (contralateral and ipsilateral) mice. (C) Brain-serum ration at 24 hours end point or after a second does (n=4 in sham group, n=3 in 24h group, n=3 in second dose group).

We calculated the average brain to serum ratio, for samples which have both measurements available. The average ratios were about 0.39 in the contralateral and 0.68 in the ipsilateral brain at 24hr. With a second dose, the brain-to-serum ratios were 0.31 and 0.30 in contralateral and ipsilateral brain, respectively (Figure 4C). These results suggest that 24 hours after stroke, Ogerin levels remain low in both serum and brain, that a second dose could help to maintain the Ogerin concentration at a higher level.

### The effect of Ogerin on ischemia-induced signaling at 24 hour after reperfusion

GPR68 activation stimulates both PKC- and PKA-dependent pathways within minutes [24]. In previous studies, we found that Ogerin sustained the PERK-eIF2a signaling during 3-6 hours after reperfusion. Here, we asked whether Ogerin alters signaling at 24 hr after reperfusion. At 24 hr after tMCAO and Ogerin IV, the contralateral side of the brain exhibited similar levels of phosphorylation of PKC and PKA substrates, whereas there was a trend of decrease in ipsilateral side (Figure 5A, B). Mammalian target of rapamycin (mTOR) plays a key role in cell survival, inflammation. We observed a decrease of p-mTOR in ipsilateral brain compared to contralateral brain (Figure 5D). The ipsilateral side of the brain exhibited decreased total CREB but with increased pCREB:CREB ratio (Figure 5C). These changes were similar in vehicle- and Ogerin-treated groups. We also notice the pJNK increased in ipsilateral brain compared to contralateral brain (Figure 5D).

**Figure 5.**
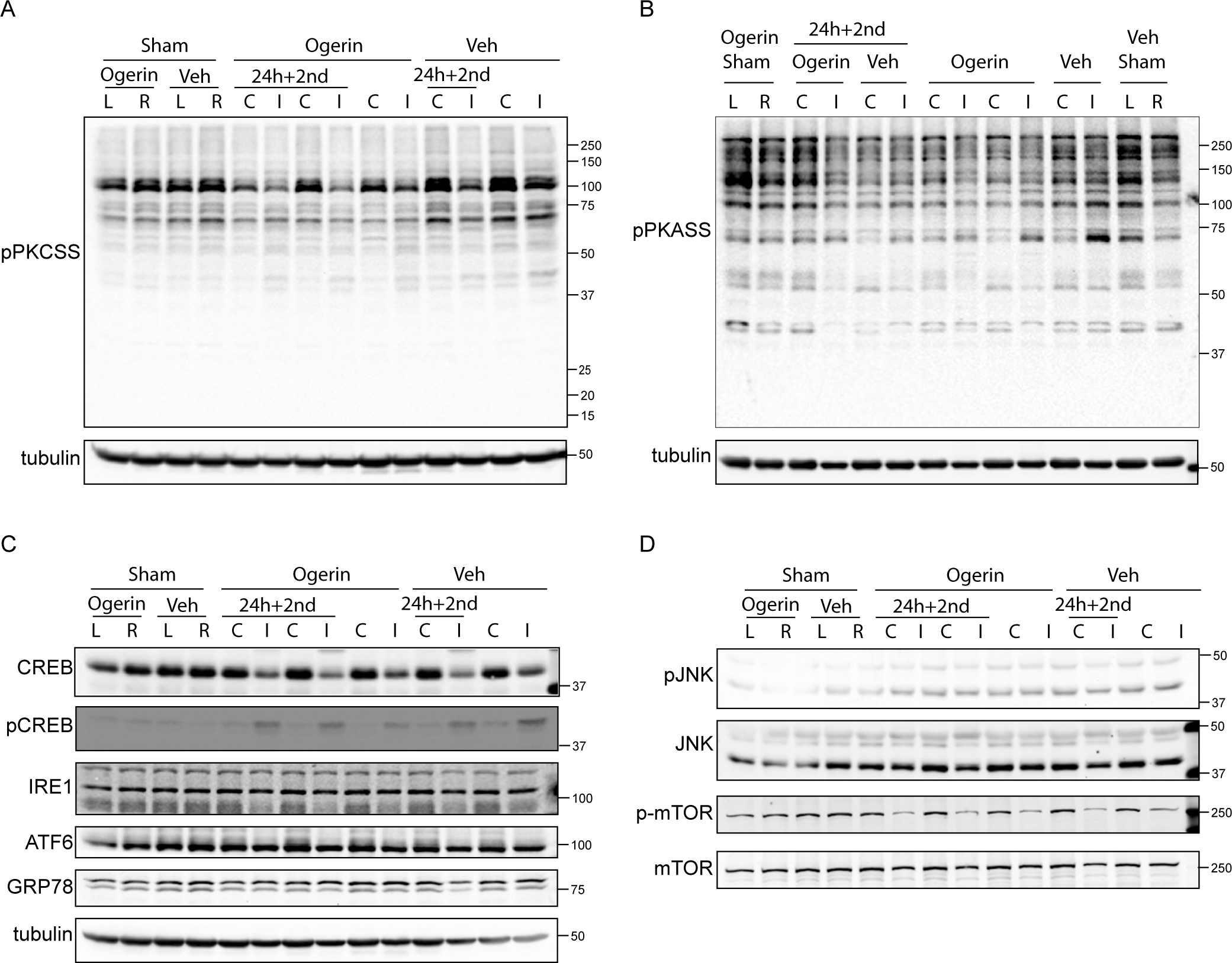
Ogerin and signaling pathways. (A-D) Representative western blot images of PKCSS (A), PKASS (B), pCREB and UPR pathways (C), p-mTOR and pJNK (D). Brain samples were collected 24 hours after tMCAO, 2mg/kg vehicle (Veh) or 2mg/kg Ogerin was delivered at reperfusion. Second dose was delivered around 20 hours after tMCAO. I: represented ipsilateral and C represented contralateral.

The unfolded protein responses (UPR) is one of important processes for ischemic tolerance [25, 26]. The three branches of UPR, ATF6, IRE1, and PERK, can exhibit different time-dependent regulations. In our previous studies, we have shown that Ogerin stimulates the PERK branch of unfolded protein responses (UPR) at 3-6 hours after reperfusion but had little effect at 24 hr [19]. Here, we examined the effect of Ogerin on ATF6 and IRE1. The result showed that there was a decreasing trend of IRE1 in ipsilateral brain compared to contralateral brain. ATF6 shows similar decreasing trend in ipsilateral brain compared to contralateral side of brain. The 78-kDa glucose-regulated protein (GRP78), also known as Bip, acts as a regulator in UPR [27]. In our results, GRP78/BIP level showed similar level in ipsilateral and contralateral brain.

## Discussion

Here, we used LC-MS analysis to determine the distribution and dynamics of Ogerin in vivo. Among all major organs, Ogerin exhibited highest accumulation in liver. The pharmacokinetic analysis in serum showed that Ogerin exhibits alpha of around 11 minute and beta of elimination of 67 min. The brain-to-serum ratio did not show significant difference in sham and tMCAO. Further, the Western blotting analysis showed that Ogerin attenuated both baseline and ischemia-induced phosphorylation of PKC substrates.

The data presented is the first PK result for Ogerin. It is worth noting that the pharmacokinetic data presented here were obtained in female animals at around 40 weeks of age. Due to the differences in metabolism and enzymatic activities [28, 29], it is possible that male and female animals can exhibit different kinetics. Age is also one important consideration. To gain more accurate projection of Ogerin level in vivo, it will be of future interest to investigate its pharmacokinetics in males and using aged mice. Nevertheless, the result here showed that Ogerin level in the serum fits well with a two-component decay. The elimination of around 67 min suggests that Ogerin level declines rapidly in serum. Consistent with this result, Ogerin in serum or brain at 24 hr was low or below detection limit. A second dose delivered around 20 hours boosted the level of Ogerin in serum and brain to ∼5 ng/ml and ∼0.7 ng/g, respectively. This finding suggests that the elimination of Ogerin is fast. For achieving long-term effect, daily injection or using an osmotic pump is needed.

In previous studies, we and others have shown that Ogerin stimulates the activation of both PKC and PKA pathways [16, 19, 30]. These studies have emphasized signaling changes within few hours. Here, we examined the signaling at 24 hours after reperfusion with Ogerin or vehicle delivered at reperfusion. It is somewhat unexpected to observe that Ogerin treatment reduced phosphorylation of PKC substrates and PKA substrates at this time point. On the other hand, the change in CREB phosphorylation was in the opposite trend as PKC substrates, with ipsilateral side showing a reduced phosphorylation of PKC substrates, decreased CREB expression but increased CREB phosphorylation. Further analysis is needed to determine the functional linkage of these signaling changes to ischemic injury.

GPR68 activation leads to protection through multiple mechanisms. In one study, activation of GPR68 reduces ferroptosis through PI3K/Akt/Nrf2 in spinal cord ischemia model [31]. We previously showed that part of the protection depends on PKC-dependent activities [14]. Ogerin, through stimulating GPR68, prolongs the activation of PERK-eIF2a branch of UPR [19]. This effect lasts until about 6 hours after reperfusion. This time window fits the projected Ogerin kinetics from this study. Together, these data suggest that the enhancement of PERK signaling correlates with the level of Ogerin in brain tissue. Besides PERK, the inositol-requiring enzyme 1 (IRE1) is a conserved ER-bound protein, also is a key sensor of ER stress and activator of the UPR [32]. The third branch of UPR was activating transcription factor 6 (ATF6) [33], is an important molecule in UPR, also plays an essential role in inflammatory and autoimmune diseases. Our previous study showed that the effect on PERK was diminished at 24 hours. However, the result here showed that IRE1 decreased with Ogerin treatment.

In summary, we presented here a first PK type analysis of Ogerin. The findings provide important information for understanding the in vivo kinetics of Ogerin and designing of more efficient treatment protocols. The Western blot result revealed new insights into the effect of Ogerin at 24 hr, on both signaling and UPR pathways.

## Acknowledgement

We thank Dr. Weqi Fu, Angela Barattini, Cory Beswick (Tulane University) for technical assistance, Dr. Wenyan Sun for surgery training, and Dr. William Gutheil and Dr. Nitish Mishra from University of Missouri-Kansas City for pilot experiments. We thank Dr. Yingjun Ma, Lucille Yu, and Ning Ren from WuXi AppTec for assistance with LC-MS analysis.

## Author contribution

Conceptualization X.Z.; Transient middle cerebral artery occlusion (tMCAO) surgery by Y.Z.; sample collection and preparation by Y.Z., S.J.; Western blotting: Y.Z.; data analysis by Y.Z.; original draft by Y.Z.; editing by X.Z.; funding acquisition X.Z. All authors have read and agreed to the final version of the manuscript.

## Data availability

All data is available in the main text and/or figures. Additional inquiry can be directed to the corresponding author.

## References

1. Feigin, V.L., et al., World Stroke Organization: Global Stroke Fact Sheet 2025. Int J Stroke, 2025. 20(2): p. 132–144.

2. de Havenon, A. and K.N. Sheth, Stroke’s Resurgence as a Leading Cause of Death in the United States. Stroke, 2025. 56(9): p. e199–e200.

3. Fan, J., et al., Global Burden, Risk Factor Analysis, and Prediction Study of Ischemic Stroke, 1990-2030. Neurology, 2023. 101(2): p. e137–e150.

4. Chung, C.P., Types of Stroke and Their Differential Diagnosis, in Primer on Cerebrovascular Diseases. 2017. p. 372–376.

5. Tran, H., Understanding ischemic strokec: auses, symptoms and treatment options. J. Experi. Stroke. Trans. Med., 2024. 16(**4**): p. 219–220.

6. Hu, H.J. and M. Song, Disrupted Ionic Homeostasis in Ischemic Stroke and New Therapeutic Targets. J Stroke Cerebrovasc Dis, 2017. 26(12): p. 2706–2719.

7. Sarkar, S., et al., Cerebral ischemic stroke: cellular fate and therapeutic opportunities. Front Biosci (Landmark Ed), 2019. 24(3): p. 435–450.

8. Zha, X.M., Z.G. Xiong, and R.P. Simon, pH and proton-sensitive receptors in brain ischemia. J Cereb Blood Flow Metab, 2022. 42(8): p. 1349–1363.

9. Wang, E., et al., Mapping tissue pH in an experimental model of acute stroke - Determination of graded regional tissue pH changes with non-invasive quantitative amide proton transfer MRI. Neuroimage, 2019. 191: p. 610–617.

10. Heo, H.Y., et al., Amide proton transfer imaging in stroke. NMR Biomed, 2023. 36(6): p. e4734.

11. Zou, C., et al., pH-sensitive probes for ischemic stroke: advancing early detection and accurate grading. Front Mol Biosci, 2026. 13: p. 1821099.

12. Halsey, J.H., et al., The role of tissue acidosis in ischaemic tissue injury: the concept of the pH integral. Neurol Res, 1988. 10(2): p. 97–104.

13. Imenez Silva, P.H. and C.A. Wagner, Physiological relevance of proton-activated GPCRs. Pflugers Arch, 2022. 474(5): p. 487–504.

14. Wang, T., et al., GPR68 Is a Neuroprotective Proton Receptor in Brain Ischemia. Stroke, 2020. 51(12): p. 3690–3700.

15. Li, X., et al., Neuroprotective effects of GPR68 against cerebral ischemia-reperfusion injury via the NF-κB/Hif-1α pathway. Brain Res Bull, 2024. 216: p. 111050.

16. Huang, X.P., et al., Allosteric ligands for the pharmacologically dark receptors GPR68 and GPR65. Nature, 2015. 527(7579): p. 477–83.

17. Yu, X., et al., *Design, Synthesis,* and Characterization of Ogerin-Based Positive Allosteric Modulators for G Protein-Coupled Receptor 68 (GPR68). J Med Chem, 2019. 62(16): p. 7557–7574.

18. Sun, W., et al., Potentiation of GPR68 alleviates post-ischemia BBB dysfunction and brain edema in mice. Exp Neurol, 2026. 403: p. 115850.

19. Sun, W., et al., Time-Dependent Potentiation of the PERK Branch of UPR by GPR68 Offers Protection in Brain Ischemia. Stroke, 2024. 55(10): p. 2510–2521.

20. Percie du Sert, N., et al., The ARRIVE guidelines 2.0: Updated guidelines for reporting animal research. PLoS Biol, 2020. 18(7): p. e3000410.

21. Friedrich, J., U. Lindauer, and A. Höllig, Procedural and Methodological Quality in Preclinical Stroke Research-A Cohort Analysis of the Rat MCAO Model Comparing Periods Before and After the Publication of STAIR/ARRIVE. Front Neurol, 2022. 13: p. 834003.

22. Longa, E.Z., et al., Reversible middle cerebral artery occlusion without craniectomy in rats. Stroke, 1989. 20(1): p. 84–91.

23. Williams, M.D., et al., pH-sensing GPR68 inhibits vascular smooth muscle cell proliferation through Rap1A. Am J Physiol Heart Circ Physiol, 2024. 327(5): p. H1210–h1229.

24. Zhou, G. and X.M. Zha, GPR68 Contributes to Persistent Acidosis-Induced Activation of AGC Kinases and Tyrosine Phosphorylation in Organotypic Hippocampal Slices. Front Neurosci, 2021. 15: p. 692217.

25. Ouyang, P.D., et al., *The mechanism of synpo in intracerebral hemorrhage using a* zebrafish *model*. Yi Chuan, 2026. 48(4): p. 407–420.

26. Sanderson, T.H., M. Gallaway, and R. Kumar, Unfolding the unfolded protein response: unique insights into brain ischemia. Int J Mol Sci, 2015. 16(4): p. 7133–42.

27. Lee, A.S., Stress-induced translocation of the endoplasmic reticulum chaperone GRP78/BiP and its impact on human disease and therapy. Proceedings of the National Academy of Sciences, 2025. 122(30): p. e2412246122.

28. Kaikaew, K., et al., Sex Difference in Corticosterone-Induced Insulin Resistance in Mice. Endocrinology, 2019. 160(10): p. 2367–2387.

29. de Souza, G.O., F. Wasinski, and J. Donato, Characterization of the metabolic differences between male and female C57BL/6 mice. Life Sciences, 2022. 301: p. 120636.

30. Bell, T.J., et al., Ogerin mediated inhibition of TGF-beta(1) induced myofibroblast differentiation is potentiated by acidic pH. PLoS One, 2022. 17(7): p. e0271608.

31. Yang, R., et al., Pharmacological Activation of GPR68 Attenuates Ferroptosis in Spinal Cord Ischemia/Reperfusion Injury Through PI3K/Akt-Mediated Nrf2 Antioxidant Pathway. Inflammation, 2025. 48(6): p. 4289–4301.

32. Spinetti, E., et al., *Structural Dynamics of IRE1 and its Interaction with Unfolded Peptides*. 2026, eLife Sciences Publications, Ltd.

33. Lei, Y., et al., Molecular mechanism of ATF6 in unfolded protein response and its role in disease. Heliyon, 2024. 10(5): p. e25937.

